# New insights into the influence of plant and microbial diversity on denitrification rates in a salt marsh

**DOI:** 10.1101/2020.08.03.234666

**Authors:** Olivia U. Mason, Patrick Chanton, Loren N. Knobbe, Julian Zaugg, Behzad Mortazavi

## Abstract

Coastal salt marshes are some of the most productive ecosystems on Earth, providing numerous services such as soil carbon storage, flood protection and nutrient filtering, several of which are mediated by the sediment microbiome associated with marsh vegetation. Here, nutrient filtering (nitrate removal through denitrification) was examined by determining microbial community structure (16S rRNA gene iTag sequencing), diversity, denitrification rates and metabolic potential (assembled metagenomic sequences) in collocated patches of *Spartina alterniflora* (*Spartina*) and *Juncus roemerianus* (*Juncus*) sediments. The iTag data showed that diversity and richness in *Spartina* and *Juncus* sediment microbial communities were highly similar. However, microbial community evenness differed significantly, with the most even communities observed in *Juncus* sediments. Further, denitrification rates were significantly higher in *Juncus* compared to *Spartina*, suggesting oscillations in microbial abundances and in particular the core microbiome identified herein, along with plant diversity influence marsh nitrogen (N) removal. Amplicon and assembled metagenome sequences pointed to a potentially important, yet unappreciated Planctomycetes role in N removal in the salt marsh. Thus, perturbations, such as sea-level rise, that can alter marsh vegetation distribution could impact microbial diversity and may ultimately influence the ecologically important ecosystem functions the marsh sediment microbiome provides.

## Introduction

As an important component of the terrestrial biological carbon pool wetlands (Dixon and Krankina, 1995; Chmura *et al.*, 2003; Fourqurean *et al.*, 2012; Duarte *et al.*, 2013) play a critical role in the global carbon cycle (Sahagian and Melack, 1998; Chmura *et al.*, 2003). Coastal wetlands at 32,000 km^2^ (Mitsch and Gosselink, 2015) are estimated to store 44.6 Tg carbon yr ^-1^ (Chmura *et al.*, 2003) acting as an important carbon sink (Adams *et al.*, 1990; Watson *et al.*, 2000; Howard *et al.*, 2017). In coastal wetlands, production rates are high even relative to agricultural land (Odum, 1959). In fact, salt marshes, and in particular Southeastern U.S. salt marshes (Pomeroy and Wiegert, 1981; Newell, 2001), are some of the most productive ecosystems on Earth (Tiner, 1984). Over half of plant biomass production is in the belowground rhizomes and roots per annum (Valiela *et al.*, 1976; Gallagher and Plumley, 1979; Schubauer and Hopkinson, 1984), which is a primary contributor to carbon sequestration in marshes (DeLaune *et al.*, 1983; Nyman *et al.*, 2006). Belowground conditions are water-saturated, which limits the diffusion of oxygen thereby slowing decomposition rates (Solomon *et al.*, 2007) and contributing to substantial carbon accumulation over time (Chmura *et al.*, 2003) and a large belowground carbon sink (Chmura *et al.*, 2003; Bridgham *et al.*, 2006; Langley and Megonigal, 2010; Mcleod *et al.*, 2011). Thus, coastal salt marshes can overlie millennial-aged deposits of organic matter (Ward *et al.*, 2008; Duarte *et al.*, 2013; Elsey-Quirk and Unger, 2018). Beyond long term soil carbon storage, flood protection and nutrient filtering are important ecosystem services provided to human communities (Mitsch and Gosselink, 2015). These services provide an underappreciated economic value of thousands of dollars per hectare per year (Costanza *et al.*, 2014).

Salt marsh vegetation is typically zonated with the more stress tolerant species in the lower elevation zones that frequently flood, while less stress tolerant species are in the higher elevation regions. Within these zones differences in porewater geochemistry (Koretsky *et al.*, 2008), carbon sequestration capacity (Elsey-Quirk, Seliskar, Sommerfield, *et al.*, 2011), and biogeochemical reactions, mediated by the microbial community, have been attributed to vegetation type (Koretsky *et al.*, 2008; Oliveira *et al.*, 2010, 2012; Davis *et al.*, 2011; Moffett and Gorelick, 2016), variable hydrology (Linthurst and Seneca, 1980; King *et al.*, 1982), as well as the marsh biota on belowground processes (Kostka *et al.*, 2002; Gribsholt *et al.*, 2003; Dollhopf *et al.*, 2005).

In the highly productive Southeastern U.S. salt marshes (Pomeroy and Wiegert, 1981; Newell, 2001) *Spartina alterniflora* Loisel and *Juncus roemerianus* Scheele (henceforth referred to as *Spartina* and *Juncus*) are the two dominant plants (Eleuterius, 1976; Wiegert *et al.*, 1983; Stout, 1984). These two plant types have several functional differences, with *Spartina* being more tolerant of flooding, anoxic soils, and high salinities (Mendelssohn and Morris, 2000) and is typically found in narrow bands at low elevation, while *Juncus* is less stress tolerant (Eleuterius, 1976; Wiegart and Freeman, 1990) and more abundant at higher elevations (Battaglia *et al.*, 2012). Observations of belowground processes have revealed that sediment geochemical characteristics indicated higher productivity and transport of greater quantities of oxygen to the subsurface in *Juncus* than in *Spartina* marshes (Koretsky *et al.*, 2008; Koop-Jakobsen and Wenzhöfer, 2015).

Of particular importance is the dynamic above-to-belowground plant-microbe interaction that leads to the removal of nitrate through denitrification by the belowground microbial community. This ecosystem function is a highly desirable service provided by marshes (Costanza *et al.*, 1997; Hopkinson and Giblin, 2008), as it leads to a reduction in nitrogen loading to the coastal ocean (Valiela *et al.*, 2000; Valiela and Cole, 2002; Velinsky *et al.*, 2017). Denitrification is recognized as the primary pathway by which nitrogen is lost from wetlands (Mitsch and Gosselink, 2015) with high denitrification rates reported in coastal marshes (Valiela and Teal, 1979; Hopkinson and Giblin, 2008). For example, the belowground microbial community has been shown to account for 84% of fixed nitrogen removal in salt marshes (Valiela and Teal, 1979; Koop-Jakobsen and Giblin, 2010), primarily via denitrification (Anderson *et al.*, 1997; Tobias *et al.*, 2001; Hamersley and Howes, 2005). Elsey-Quirk *et al.* (2011) reported 46% higher N turnover in *Juncus* than in *Spartina*, highlighting the importance of plant species on belowground processes.

The microbial communities in salt marsh sediments are remarkably diverse (Bowen *et al.*, 2012), and environmental heterogeneity resulting from the vegetation or the hydrology of the marsh impacts the structure of these microbial communities. The marsh vegetation in each zone provides unique substrates in their rhizosphere (Oliveira *et al.*, 2012) that can favor some members of belowground microbial community over others (Cleary *et al.*, 2016; Rietl *et al.*, 2016). Further, it was recently reported that plant lineage rather than environmental conditions structures bacterial communities (Bowen *et al.*, 2017), suggesting a tight coupling between plants and the associated sediment microbiome. Additionally, the different microbial communities within marsh vegetation zones can result in varying biogeochemical rates. For example, Oliveira et al. (2010) found different bacterial communities in two monospecific stands of marsh grasses and attributed the variable activity of the heterotrophic community to the influence of the marsh plants on belowground processes. However, because plant zonation in marshes mainly occurs along an elevation gradient, separating the impact of the vegetation as opposed to that of the elevation (Cao *et al.*, 2008), which influences the inundation and drainage of the marsh, remains challenging.

Thus, our study site provided a unique opportunity to sample sediments from *Spartina* and *Juncus* patches collocated at the same elevation in a Gulf of Mexico salt marsh. This mixed marsh allowed us to test the hypothesis that distinct microbial communities would be associated with the two plant types despite being exposed to similar inundation cycles and that each plant type’s unique microbial community would translate to different denitrification rates. To test our hypothesis we used 16S rRNA gene iTag sequencing (ASVs) which enabled us to uncover differences in microbial community composition and diversity in sediments dominated by these two marsh plants with high resolution. Furthermore, we determined how different microbial structure and diversity within these two plant types might influence denitrification rates, which were quantified in the two different plants. Finally, we assembled metagenomic sequence data to determine the potential function of indigenous salt marsh microbes. Collectively this approach allowed us to test our primary hypothesis regarding plant-microbe interactions, microbial community structure, diversity and function.

## Materials and Methods

### Study site

The salt marsh on Dauphin Island (30°15.43’N, 88°07.438’W) is dominated by *Spartina* and is interspersed with *Juncus*. Tides are diurnal with mean tidal amplitude of less than 0.5 m. The marsh is flooded on every high tide. Elevation of *Juncus* and *Spartina* patches were measured with a Trimble Real Time Kinematic (RTK) GPS (TSC-2 controller and Trimble-R8 Model-3 rover) on five separate occasions and did not differ (difference: 0.01 ± 0.02 m). Midday air temperatures at the site ranges from 9 to 31°C and salinity ranges from 12 to 32 ppt (Wilson *et al.*, 2015).

### Sample collection and processing

#### Denitrification rates

Replicate sediment cores (95 mm ID; n=5) were collected from *Spartina* and *Juncus* dominated sediments during the April 2015 to October 2016 period. Each core was sectioned between the 0-2 and 5-7 intervals and homogenized. Denitrification rates were measured following the acetylene inhibition technique (Sorensen, 1978), similar to the procedures used by Dollhopf et al. (2005). Twenty grams of sediments from each core and each depth and filtered site water were added to serum vials and amended with nitrate to a concentration of 100 µM. Samples were sealed with a butyl rubber stopper, capped and flushed with N_2_ gas for 10 min. After the addition of C_2_H_2_ (10 % v/v) and a 1-h incubation, headspace gas samples were injected into evacuated 12 mL Exetainer vials and N_2_O production was quantified with a Shimadzu GC-2014 with an electron capture detector (GC-ECD) within 24 h. It is recognized that the acetylene inhibition technique inhibits nitrification and can lead to incomplete inhibition of N_2_O reduction (Seitzinger et al. 1993). To minimize these impacts the incubation time was held to 1 hr.

#### Porewater nutrient concentrations

Porewater nutrient concentrations were periodically collected from triplicate sippers placed in *Juncus* and *Spartina* dominated patches. Sippers were equipped with a 5-cm sampling window (centered at 10 cm depth) of porous plastic (Porex, 25-40 µm pore size) (Neubauer, 2013). Prior to sampling, the sippers were purged of water and flushed with N_2_ gas to maintain anoxia. Samples were then withdrawn with a syringe, filtered through GF/F filters, kept on ice and returned to the lab where they remained frozen until analysis. Nitrate, ammonium and phosphate concentrations were analyzed with standard wet chemical techniques modified for the Skalar SAN+ Autoanalyzer.

#### Sediment hydrogen sulfide profiles

In May and August 2015 additional sediment cores were collected from the *Spartina* and *Juncus* patches and brought back to the laboratory for profiling of the porewater hydrogen sulfide concentrations. Cores were kept at *in situ* temperature and submerged in site water that was maintained normoxic. Hydrogen sulfide (measured as HS^-^) concentrations were determined with a Unisense microelectrode system with H_2_S (H_2_S-500) sensor calibrated according to manufacturer’s instructions. Concentrations were recorded at the sediment–water interface to a depth of 20 mm at 1 mm intervals with a micromanipulator in each profile.

#### DNA extraction and purification

DNA was extracted from 1 g of sediment from 194 samples, which includes 5 replicates per sample from *Spartina* and *Juncus* dominated sediments that were collected April 2015 to October 2016, using a modified CTAB extraction buffer ((10% CTAB (hexadecyltrimethylammonium bromide), 1M NaCl and 0.5M phosphate buffer, pH 8) with 0.1M ammonium aluminum sulfate, 25:24:1 phenol:chloroform:isoamyl alcohol) and subjecting it to bead beating using a FastPrep-24 (MP Biomedicals, Solon, OH) following the protocol as described by Gillies et al. (2015) with the following modification: the first and second extractions were combined after the ethanol wash with 50 ul EB buffer to maximize DNA yields. DNA was purified using the QIAGEN AllPrep DNA/RNA Kit (QIAGEN, Germantown, MD) following the manufacturer’s protocol.

#### 16S rRNA gene sequencing and analysis

The microbial community structure in all samples was analyzed using iTag sequencing. 16S rRNA genes were amplified from 10 ng of purified genomic DNA in duplicate using the modified archaeal and bacterial primers 515FB and 806RB (Apprill *et al.*, 2015; Parada *et al.*, 2015) in accordance with the protocol described by Caporaso et al. (2011, 2012) and used by the Earth Microbiome Project (http://www.earthmicrobiome.org/emp-standard-protocols/16s/), with a slight modification: the annealing temperature used was 60 °C. Amplicons were sequenced using an Illumina MiSeq in 250 × 250 bp mode. These sequences will be available in the National Center for Biotechnology Information (NCBI) SRA and on the Mason server at http://mason.eoas.fsu.edu. Raw sequences were demultiplexed using QIIME2 (Caporaso *et al.*, 2010; ver. 2018.6). Demultiplexed reads were quality filtered, including chimera removal, and joined using DADA2 (Callahan *et al.*, 2016) using default parameters. The resulting exact amplicon sequence variant (ASV) (Callahan *et al.*, 2017) data was normalized using cumulative sum scaling (CSS) (Paulson *et al.*, 2013). ASV taxonomy was assigned with the SILVA database (Yilmaz *et al.*, 2014; ver. 132) in QIIME2 using classify-sklearn. To determine closest cultured representatives to specific ASVs NCBI’s 16S ribosomal RNA sequences for Bacteria and Archaea database and the Joint Genome Institute’s (JGI) nucleotide blast in IMG (single amplified genomes and metagenome assembled genomes were also searched) were used.

#### Statistical analyses

ASV data was normalized using multiple rarefaction, which was then used to determine Shannon diversity (Shannon and Weaver, 1949), richness (Chao1) (Chao, 1984) and evenness (Pielou e) (Pielou, 1966) using QIIME 2. The Shapiro-Wilk normality test was used to determine whether diversity data were normally distributed. The normalized ASV table generated using DADA2 in QIIME 2 was analyzed using non-metric multidimensional (NMDS) scaling in R using the metaMDS command in the vegan package. P-values for statistical significance of environmental variables and alpha diversity in relationship to NMDS axes were derived from 999 permutations of the data. Spearman’s rank correlation coefficients were determined in R using the psych package (Revelle, 2018). Bonferroni corrections for multiple tests were applied to p-values from Spearman’s correlations. To test for significant differences in alpha diversity metrics and normalized ASV abundances between plant type and depth Wilcox test with the Benjamini-Hochberg (B-H) and, when appropriate Kruskal-Wallis rank sum test with the B-H correction. For the statistical analyses the correction for multiple tests was based on the number of tests, with fewer tests (e.g. Spearman’s correlations) corrected with Bonferroni (Bonf.) (most conservative) and large number of tests (e.g. 32,863 ASVs) corrected with B-H.

#### Metagenomic sequencing, assembly and annotation

For shotgun metagenomic sequencing DNA extracted from sediments in *Juncus* and *Spartina* samples were sequenced by JGI using the Illumina NovaSeq Regular 270 bp fragment mode. Raw reads were processed with Trimmomatic (ver. 0.36, default settings) for adaptor removal and quality filtering. Reads were then assembled using MetaSPAdes (ver. 3.13) with default parameters. Contigs whose length was less than 500bp were removed using BBMap (ver. 38.41, https://sourceforge.net/projects/bbmap/). Quality controlled reads for each sample were mapped onto their respective assemblies using CoverM ‘make’ (ver 0.2.0, B. Woodcroft, unpublished, https://github.com/wwood/CoverM). Low quality mappings were removed with CoverM ‘filter’ (minimum identity 95% and minimum aligned length of 50%). Scaffolds for each sample were binned by providing each sample’s contigs and BAM files as input to UniteM (ver. 0.0.15, D. Parks, unpublished, https://github.com/dparks1134/UniteM) and using a minimum contig length of 1,500bp and Maxbin (ver. 2.2.4) (Wu *et al.*, 2014), MetaBAT (ver. 0.32.5) (Kang *et al.*, 2015) and MetaBAT2 (ver. 2.12.1) (Kang *et al.*, 2019) binning methods (max40, max107, mb2, mb_verysensitive, mb_sensitive, mb_specific, mb_veryspecific and mb_superspecific). Bin completeness and contamination was evaluated using CheckM (ver. 1.0.12) (Parks *et al.*, 2015). Taxonomies were assigned to each bin using GTDB-Tk (ver. 1.0.2) (Chaumeil *et al.*, 2019), which assigns an objective taxonomic classification of bacterial and archaeal genomes by placing them into domain-specific, concatenated protein reference trees provided by the GTDB (Parks *et al.*, 2018). One high quality Planctomycetes genome was obtained (bin 134) from a *Spartina* sediment sample collected in May 2015 from 0-2 cm. Bin 134 was deposited in JGI’s IMG for annotation (Chen *et al.*, 2019) and will be made publicly available in IMG. In addition to IMG annotations, blastn against the Silva database (ver. 132) was used to search for 16S rRNA genes in the metagenome assembled genome (MAG).

## Results

### Denitrification rates and nitrate, ammonium, phosphate and hydrogen sulfide concentrations

Denitrification rates differed between plant species types across all dates and depths (ANOVA, p≤0.05) and were nearly 3X higher in *Juncus* (48.0 ± 74.5 µmol N m^-2^ h^-1^) compared to *Spartina* (16.1 ± 48.0 µmol N m^-2^ h^-1^) sediments (Figure 1). Temporal patterns in denitrification differed between depths (ANOVA, depth x date, p-value ≤ 0.05). Denitrification in surficial (0-2) cm sediments decreased during the summer for both 2015 and 2016, with the lowest rates occurring in June 2015 and July and August 2016 (Figure 1A). Similarly, denitrification rates in sub-surface (5-7 cm) sediments were lowest in July and August 2016 (Figure 1B), but they were only significantly lower than 2015 rates. Although there is a trend of surficial denitrification rates being higher than sub-surface rates, the effect of depth was only significant in February 2016 (2.8-fold difference; Tukey’s HDS p=≤ 0.05) and October 2016 (2.2-fold difference; Tukey’s HSD p=≤ 0.05). In this same marsh system, porewater nitrate concentrations were similar in the *Juncus* and *Spartina* patches (3.3 ± 0.6 and 3.8 ± 0.8 µmols L^-1^, respectively) as were the ammonium concentration (56.2 ± 10.2 and 65.1 ± 16.8 µmols L^-1^ for *Juncus* and *Spartina*, respectively) (Figure 1C-D). In contrast, porewater phosphate concentrations in the *Juncus* patches (45.1 ± 9.2 µmols L^-1^) were three times higher than those in the *Spartina* patches (14.8 ± 4.1 µmols L^-1^) (Figure 1E). Porewater hydrogen sulfide concentrations profiled from 0-2 cm in May and August 2015 showed higher concentrations in *Spartina* patches compared *Juncus* patches (Figure 1, inset).

**Figure 1.**
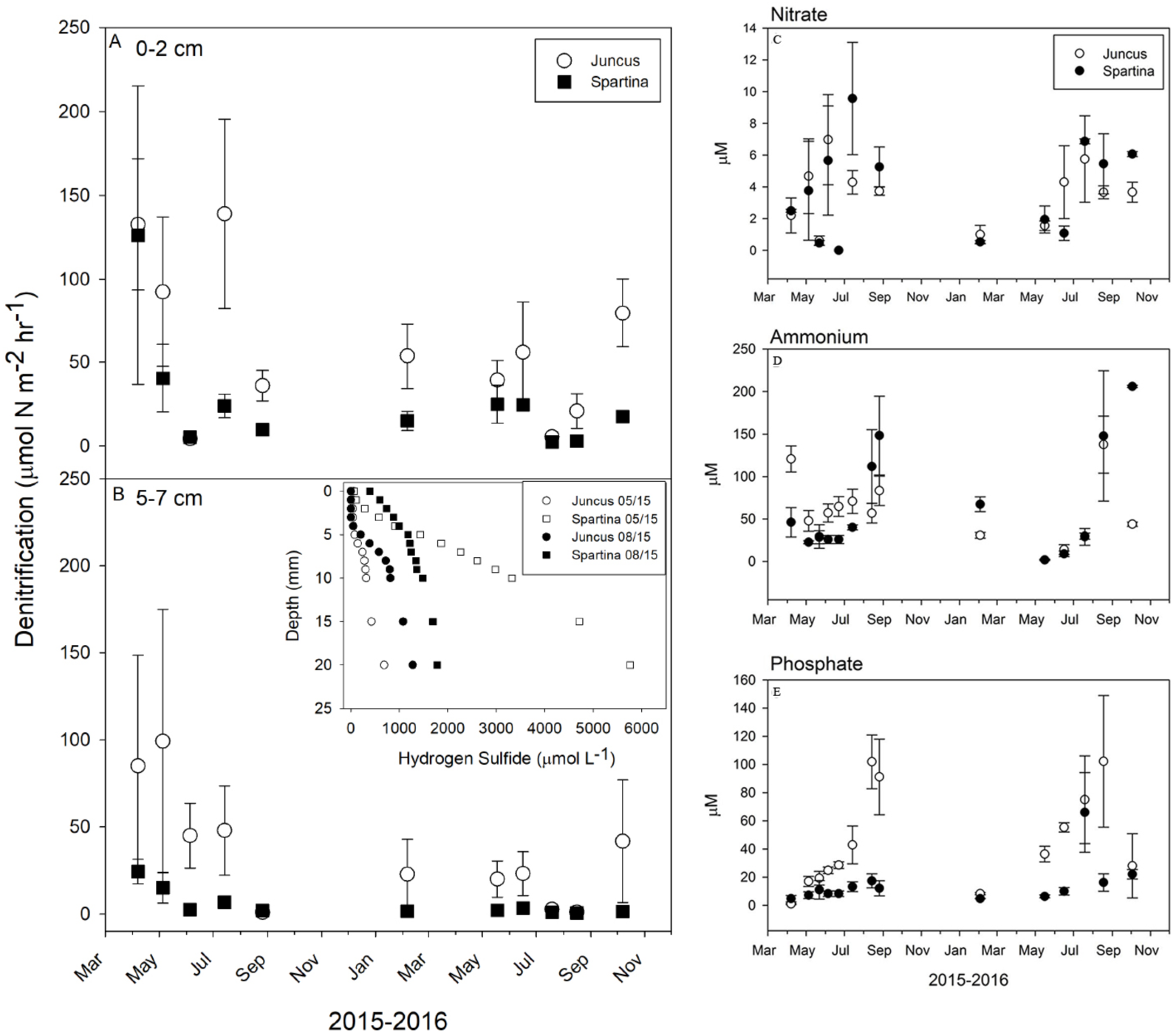
Denitrification rates from patches of *Juncus* and *Spartina* sediments from **A** 0-2 cm and **B** 5-7 cm depths. Figure 1 **C-E** shows nitrate, ammonium and phosphate concentrations. The inset shows hydrogen sulfide concentrations measured at 1 mm interval to a depth of 2 cm in May and August 2015.

### Microbial community structure in the Juncus and Spartina rhizosphere

Analysis of the normalized (CSS) 16S rRNA gene ASV data (32,863 ASVs in total) revealed that Anaerolineae in the Chloroflexi was the most abundant class in the rhizosphere of both *Juncus* and *Spartina* and at both depths, comprising more than 40% of the microbial community in some samples (Figure 2; for the purposes of discussing abundances CSS normalized data was converted to relative proportions). The relative abundance of Anaerolineae was highest in the 5-7 cm depth interval in both plants, composing 22% of the *Juncus* community and 25% of the *Spartina* community (±8% standard deviation (sd) for *Juncus* and ± 9% for *Spartina*) (Figure 2). In the 0-2 cm interval Anaerolineae relative abundances were 19% (± 6% sd for both plants) compared to the deeper horizon with 26% in *Juncus* and 32% in *Spartina* (7% sd for both). The other abundant classes were Gamma-Delta-, and Alphaproteobacteria, Bacteroidia in the Bacteroidetes, Planctomycetes; Planctomycetia and Caldithrixae, which while prevalent in all samples were more abundant in 0-2 cm in both plants (Figure 2). Thermoplasmata in the Euryarchaeota and Bathyarchaeota (formerly Miscellaneous Crenarchaeota group (Meng *et al.*, 2014)) were also abundant, but were higher in the deeper 5-7 cm interval (Figure 2).

**Figure 2.**
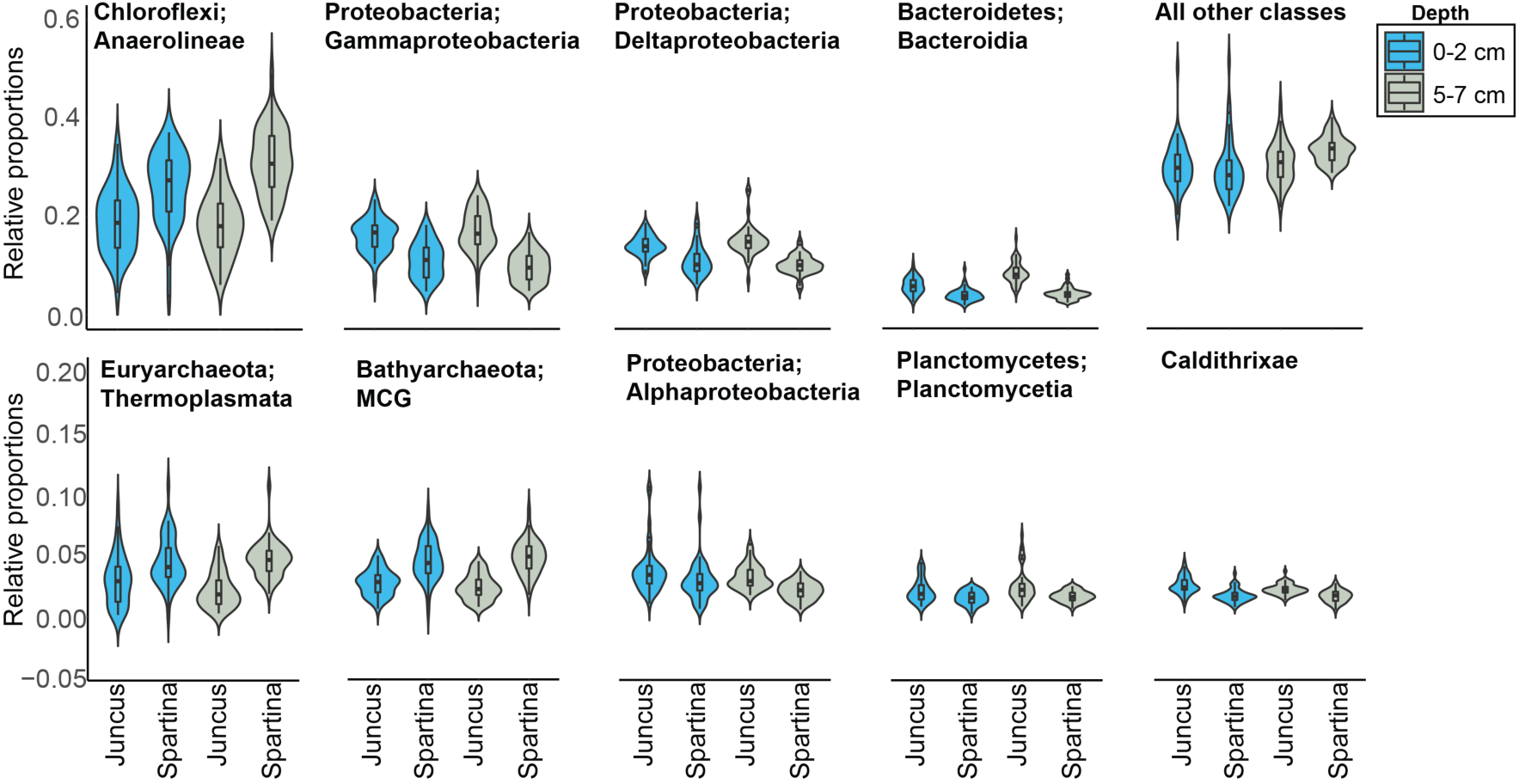
Violin plots of normalized and relativized 16S rRNA gene ASV data of the most abundant microbial classes in *Juncus* and *Spartina* sediments from 0-2 cm and 5-7 cm depths.

Spearman’s rank correlation coefficients were determined between the abundant classes (this data was CSS normalized, but not converted to relative proportions) and denitrification rates. Anaerolineae was significantly inversely correlated (Spearman’s correlation = −0.38; Bonf. corrected p-value ≤ 0.05) with denitrification rates, while Gammaproteobacteria was positively correlated (0.32), followed by Alphaproteobacteria (0.22) and Caldithrixae (0.25), with all correlations having Bonf. corrected p-values ≤ 0.05. The Planctomycetes phylum Spearman’s correlation with denitrification rates was 0.16, which was not longer significant after correcting for multiple tests.

### Microbial diversity in Juncus and Spartina sediments

Alpha diversity analysis of normalized ASV data revealed that neither diversity (Shannon) nor richness (Chao1) significantly differed between plant types and depths (Wilcoxon rank sum test; Figure 3). However, evenness (Pielou e) was significantly different between plant types (Wilcoxon rank sum test; Bonf. corrected p-value ≤ 0.05), with higher evenness values in *Juncus* than in *Spartina* (Figure 3) dominated sediments. In both plants and at both depths the 0-2 cm microbial communities in *Juncus* patches were the most even (average evenness value for this depth interval in *Juncus* was 0.92, with 1.0 being a completely even community) (Figure 3 and 4). While not significantly different, Shannon was higher in *Juncus* compared to *Spartina* patches, while Chao1 was higher in *Spartina* patches (Figure 3). Further, denitrification rates were significantly correlated with evenness (Spearman’s correlation = 0.3, Bonf. corrected p-value = ≤ 0.05). Neither diversity nor richness were significantly correlated with denitrification rates.

**Figure 3.**
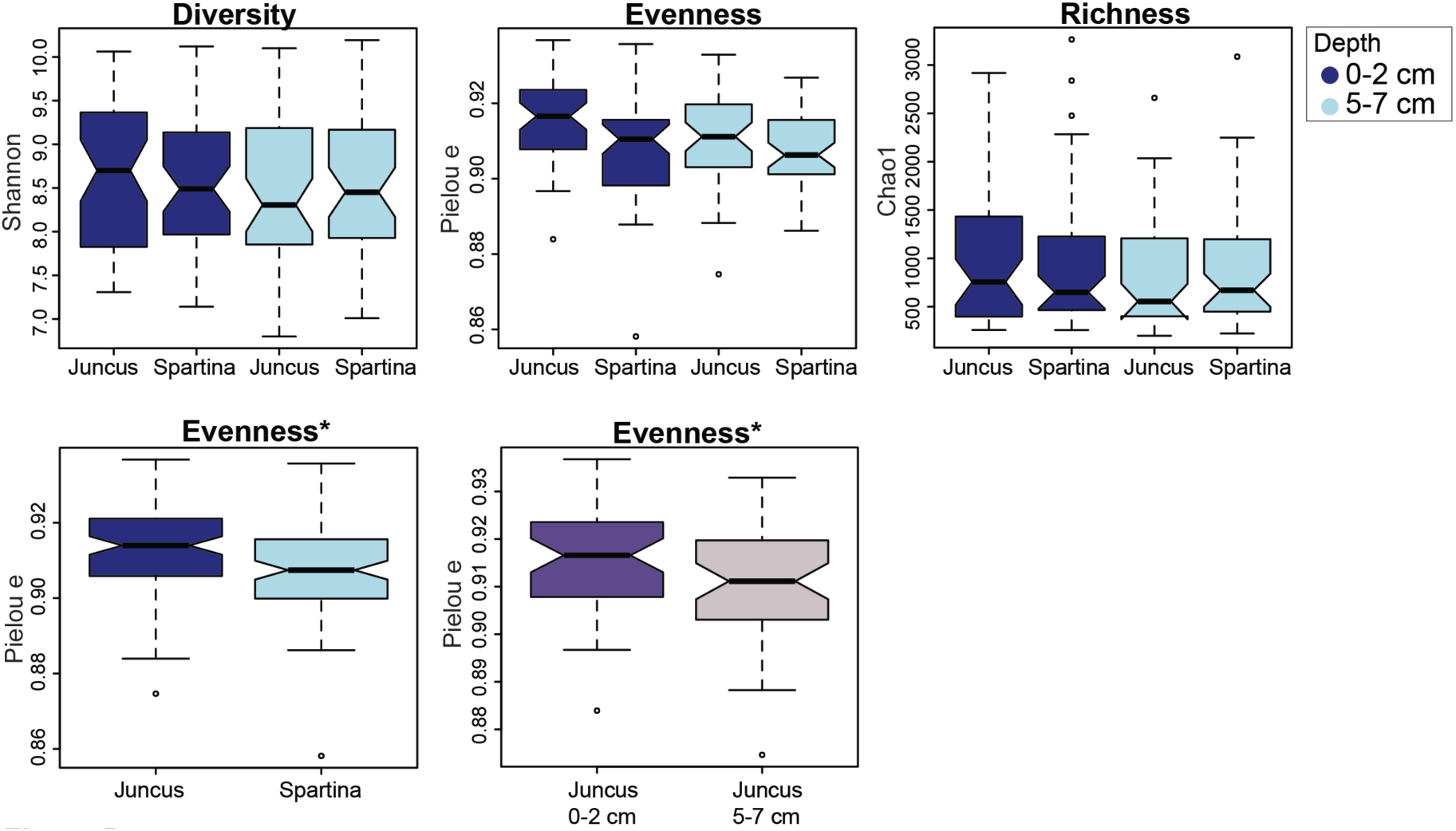
Boxplots of normalized microbial diversity metrics (Shannon), evenness (Pielou e) and richness (Chao1) calculated from 16S rRNA gene ASV data from *Juncus* and *Spartina* sediments collected at 0-2 cm and 5-7 cm depths. The asterisk indicates statistically significant differences (p-value < 0.05) in the metric evaluated.

**Figure 4.**
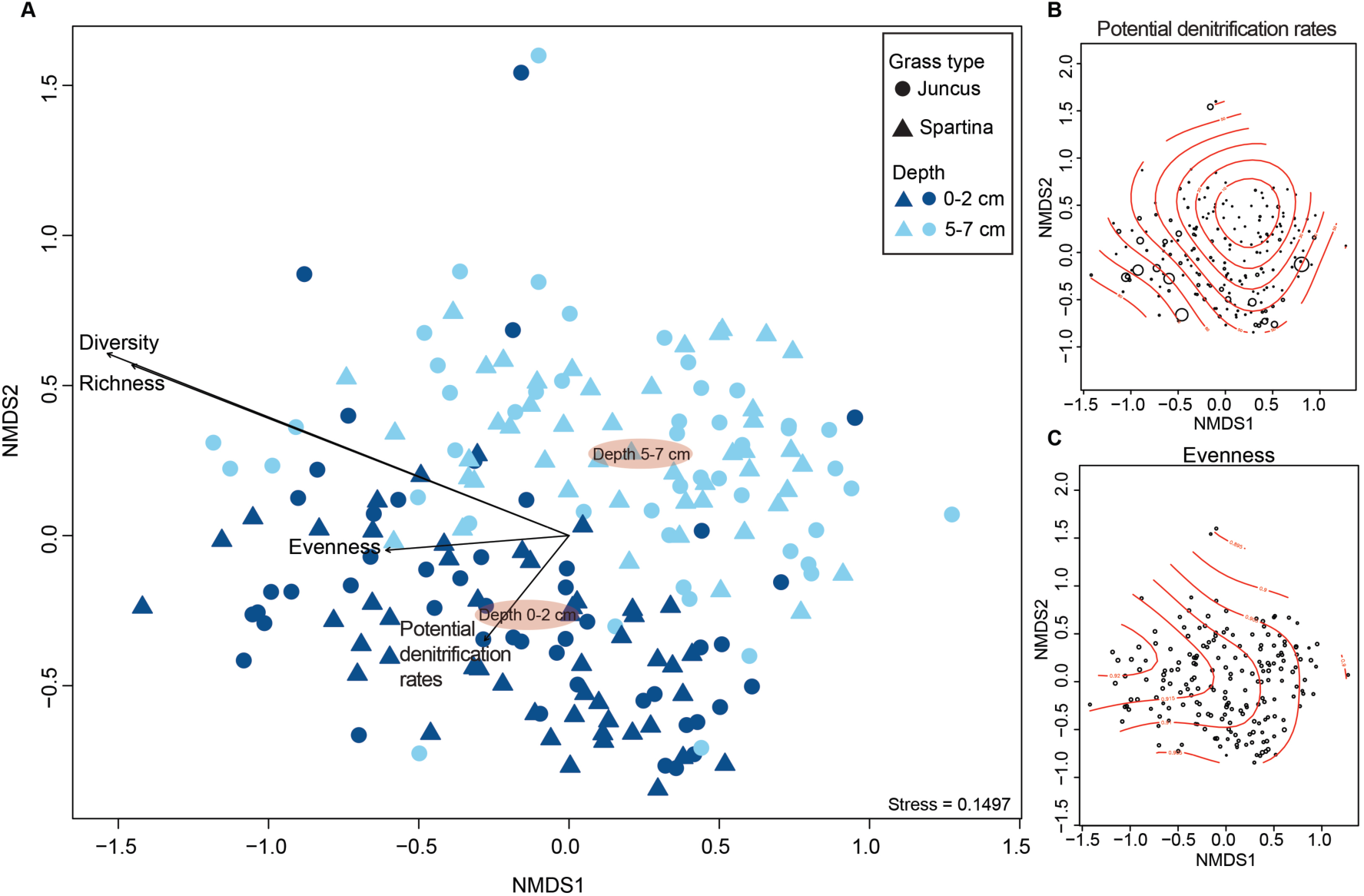
Nonmetric multidimensional scaling ordination of normalized 16S rRNA gene ASV data. **A** shows the ordination with vectors overlain and with depth centroids. **B** shows the same ordination as A but denitrification rates are shown by bubble size, with higher rates having larger bubbles. **C** shows the same ordination as A but evenness values are shown by bubble size.

### Microbial diversity, the core community and potential metabolic function

CSS normalized ASV data was analyzed using nonmetric multidimensional scaling ordination revealing that microbial communities in *Juncus* and *Spartina* had some clustering by depth, as shown by depth centroids on Figure 4. Denitrification rates and diversity metrics were overlayed onto the ordination revealing that while diversity and richness were significantly correlated with ordination axes, these vectors did not suggest a correlation with a particular plant type or depth and did not appear to be related to denitrification rates (Figure 4), consistent with the results discussed above. Similarly, evenness was significantly correlated with the ordination; however, this vector did appear to have a relationship with denitrification rates (Figure 4), which were highest in *Juncus* and *Juncus* 0-2 cm in particular.

To compare ASVs by plant type and depth further analysis was carried out on normalized ASV data using the Wilcox test with the Benjamini-Hochberg (B-H) correction for multiple tests. This comparison revealed that the normalized abundance of several ASVs (32,863 total) were significantly different when comparing 0-2 cm to 5-7 cm (both plants; 184 ASVs) and when comparing plants (both depths; 97 ASVs). A total of 56 ASVs were significantly different when comparing year, and 120 when comparing month (Kruskal-Wallis rank sum test with the B-H correction). Additionally, denitrification rates were partitioned into quartiles that represented low to median to high rates. With these categorical groupings normalized ASV abundances were tested for significant differences using the Kruskal-Wallis rank sum test with the B-H correction. A total of 119 ASVs were significantly different when comparing rates. This statistical analysis revealed a pattern - with increasing denitrification rates Planctomycetes; Planctomycetaceae abundances increased, while those classified as Chloroflexi; Anaerolineae decreased in abundance.

To determine if particular ASVs were present in all samples, suggesting a core rhizosphere microbiome associated with *Juncus* and *Spartina*, an analysis of core ASVs was carried out. This analysis revealed that no ASVs were present in all samples, but that of the 32,863 total ASVs in both plant types and in both depths, 20 were present in 95% of the samples and 21 were present in 90% of the samples. These core plant associated microbes were also among the top most abundant ASVs in all samples and their abundances across denitrification rate quartiles were significantly different (Kruskal-Wallis rank sum test with the B-H correction). Of the core ASVs eight were significantly inversely correlated with denitrification rates, including 2/3 core Anaerolineaceae representatives (Figure 5); the most abundant taxonomic group in both plant types and depths (Figure 2). However, of these core ASVs only a single one, identified as Planctomycetaceae, was significantly positively correlated with denitrification rates (Spearman’s correlation = 0.5, Bonf. corrected p-value ≤ 0.05; Figure 5). This Planctomycetes was most similar to the cultured *Planctomyces brasiliensis* (Schlesner, 1989), now referred to as *Rubinisphaera brasiliensis*, a known nitrate reducing microorganism (Schlesner, 1989; Scheuner *et al.*, 2014). This is one of the two Planctomycetes ASVs that was significantly more abundant at high denitrification rates that was discussed above.

**Figure 5.**
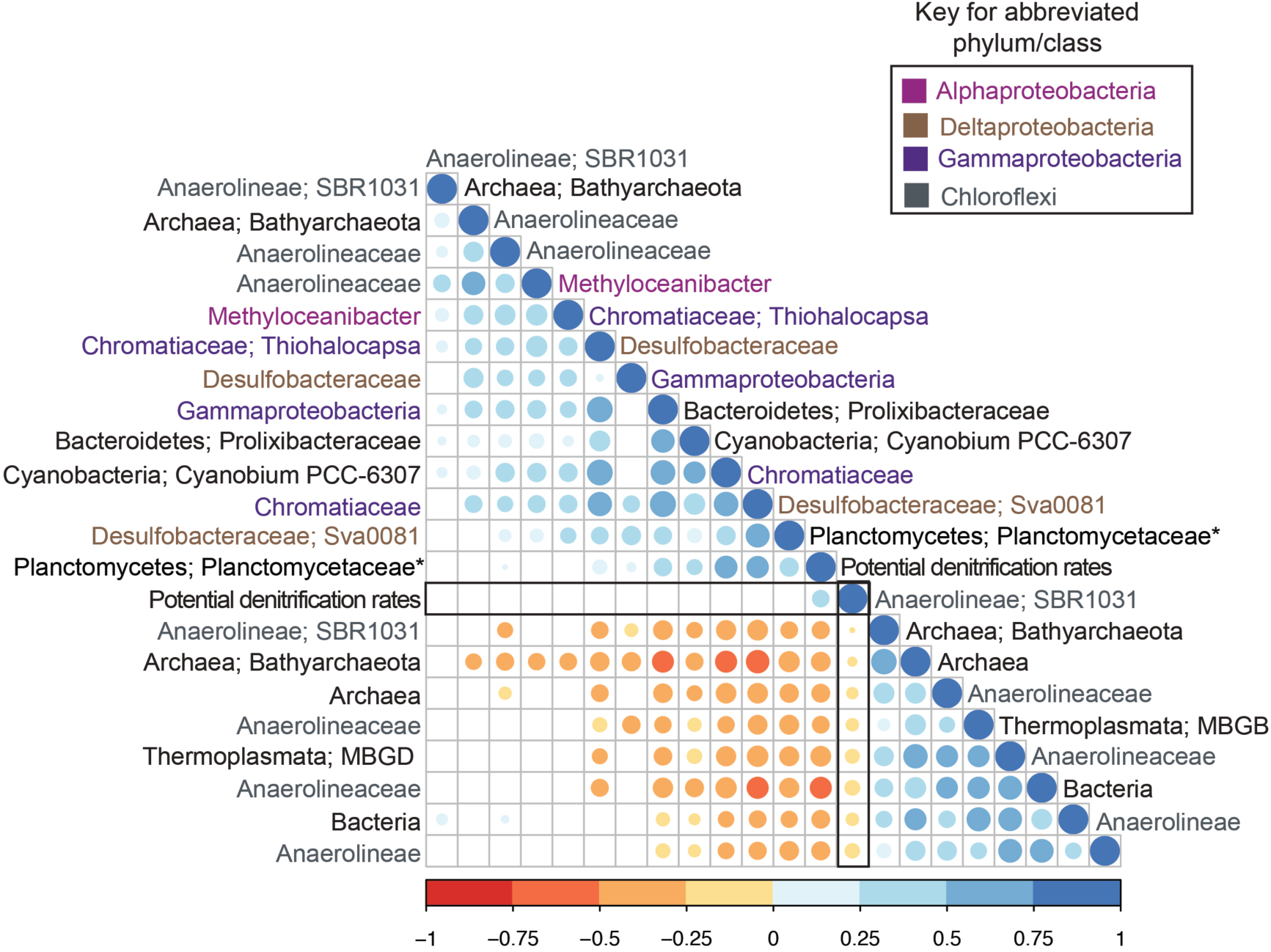
Correlogram of Spearman’s rank correlation coefficients for the core community normalized ASV abundances and denitrification rates. All core community members are a unique ASV (e.g. each Anaerolineaceae are a unique ASV), with the most finely resolved taxonomy shown. An asterisk indicates the Planctomycetes; Planctomycetaceae ASV that was significantly positively correlated with denitrification rates. Only significant correlations are shown.

The possible metabolic role of Planctomycetes in denitrification was determined by assembling metagenome data and genome annotation. From this effort, one MAG, bin 134, was obtained with a genome size of 5,867,367 bp at 68% completeness (9% contamination), with 61% GC content. Taxonomic analysis of bin 134 using GTDB-tk placed it in the Planctomycetota; Planctomycetes; *Pirellulales*; *Pirellulaceae*. There are 39 *Pirellulaceae* genomes in IMG/MER with annotations and were included in the analysis with bin 134. Genome annotation of bin 134 in IMG revealed that of the 6480 genes 6458 coded for proteins, with 3641 having functional predictions. No 16S rRNA genes were assembled, thus the MAG couldn’t be compared to the core Planctomycetes that was significantly, positively correlated to denitrification rates discussed above. The lack of 16S rRNA genes in the IMG annotation was further confirmed outside of IMG by blastn against the Silva database ver. 132. Bin 134 had the requisite genetic machinery to carry out several nitrogen respiring reactions (Figure 6). These genes are nitrate reductase ferredoxin-type protein (K02573; also identified in 4/39 *Pirellulaceae* genomes in IMG), nitrite reductase (NADH) small subunit (K00363; 38/39 *Pirellulaceae* genomes), hydroxylamine reductase (K05601; 0/39 *Pirellulaceae* genomes), nitrous oxide reductase (partial gene with no stop codon assigned to COG4263; 1/39 *Pirellulaceae* genomes), and nitrous oxidase accessory protein (K07218; 7/39 *Pirellulaceae* genomes) (Figure 6). Nitrate/nitrite transporters were also encoded in the MAG and *Pirellulaceae*. In IMG a search of bin 134’s genome for the closest homologs in all Planctomycetes revealed that uncultured *Rhodopirellula* MAGs from the *Tara* Oceans samples (Sunagawa *et al.*, 2015) encoded the most similar proteins as well as genomes from cultured *Rhodopirellula*, such as *R. baltica* (Glöckner *et al.*, 2003)*, R. europaea* (GenBank Acc. ANOF0100000 and ANMO01000001), or *Rubinisphaera brasiliensis* (GenBank Acc. NC_015174) (bit scores for all were > 100-600). The only exception was for Bin 134’s hydroxylamine reductase which was most similar to *Thermogutta terrifontis*’s hydroxylamine reductase, hybrid-cluster protein (blastp bit score was 339 with 52% similarity) (Figure 6).

**Figure 6.**
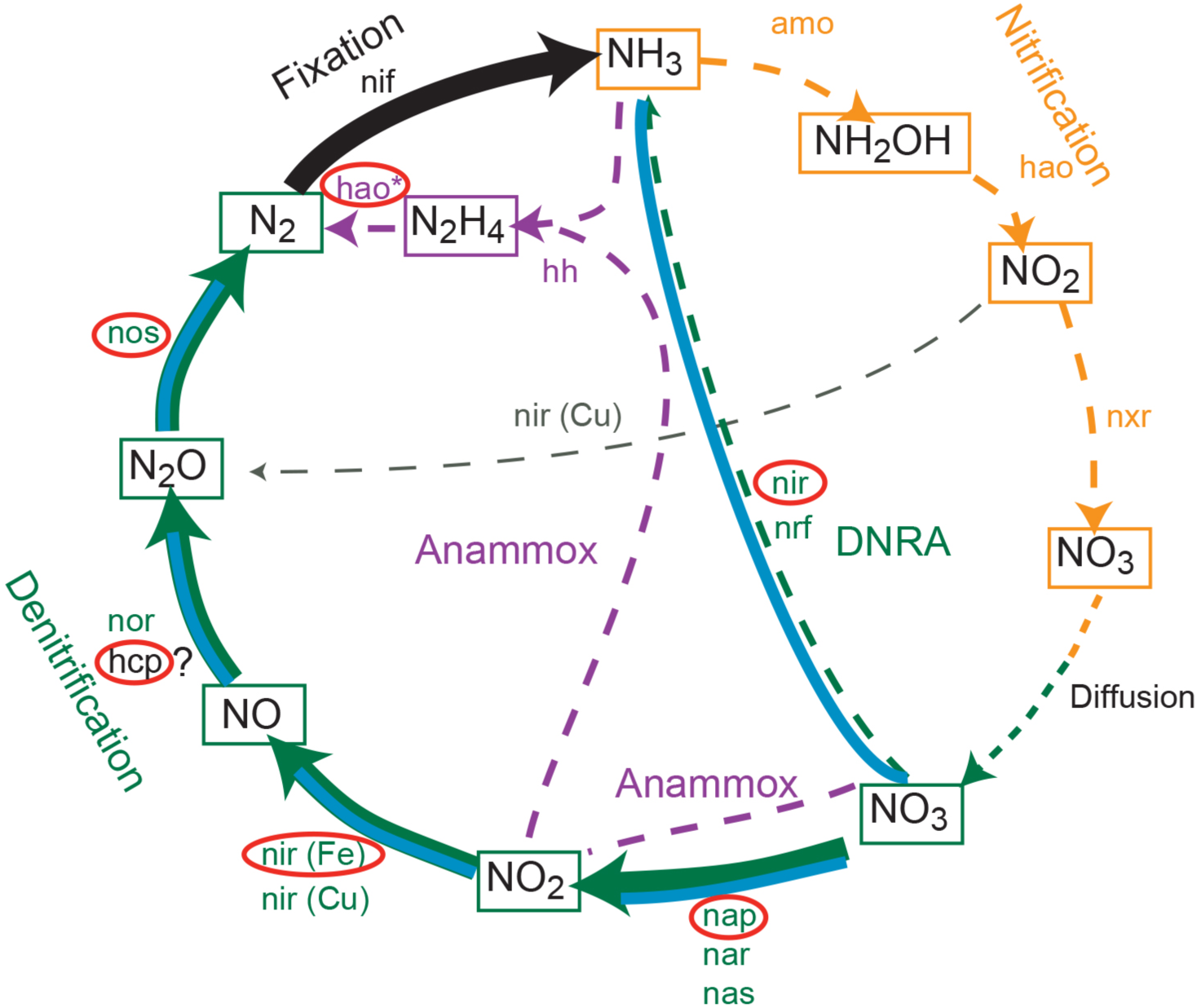
The nitrogen cycle is shown, with important pathways in salt marshes indicated with a blue line along with the pathway. Red ovals indicate genes that were identified in the Planctomycetes MAG. The role of the hcp gene in denitrification is hypothesized herein, which is indicated with a question mark following the gene abbreviation and also with an asterisk following the hao gene abbreviation.

## Discussion

Here, we collected sediment samples at a mixed marsh in the Gulf of Mexico populated by *Spartina* and *Juncus* that were at similar elevations. Sampling the different marsh plants that are exposed to similar inundation cycles allowed us to test our hypothesis that distinct microbial communities are associated with the two plant types, which influences denitrification rates in sediments from the collocated patches of *Spartina* and *Juncus*.

Examining coastal salt marsh sediment communities at higher taxonomic levels than ASVs provides the opportunity to compare our results with other studies to discern whether marshes have particular microbial classes that are regularly observed in these environments. Our analysis revealed that at the class and ASV level, the microbiome in *Spartina* and *Juncus* sediments were highly similar at both depths; therefore, we rejected our hypothesis, accepting the null hypothesis. The highly similar *Spartina* and *Juncus* microbiome was composed largely of phyla and classes that have been previously reported to be abundant in marshes. For example, Proteobacteria are regularly shown to be the most abundant phylum (Burke *et al.*, 2002; Bowen *et al.*, 2012; M. Wang *et al.*, 2016; Rietl *et al.*, 2016; Cleary *et al.*, 2017; Angell *et al.*, 2018; Angermeyer *et al.*, 2018). Specifically, in our sediment samples Gamma- and Delta- and to a lesser extent Alphaproteobacteria, were abundant. Similarly, Chloroflexi have been repeatedly shown to be abundant in marshes (Bowen *et al.*, 2012; M. Wang *et al.*, 2016; Cleary *et al.*, 2017; Angell *et al.*, 2018; Angermeyer *et al.*, 2018). In our present study, Bacteroidetes and Planctomycetes were also abundant, although less so than Proteobacteria and Chloroflexi. Bacteroidetes have been reported to be abundant in several studies (Bowen *et al.*, 2012; Cleary *et al.*, 2017), as have Planctomycetes (Bowen *et al.*, 2012), but both are typically at lower abundances relative to Proteobacteria and Chloroflexi. Thus, patterns in the salt marsh microbiome have begun to emerge, with certain microbial classes regularly observed in this environment and presumably being important in marsh function.

In assessing microbial diversity, including the individual components of diversity, richness and evenness, in sediment samples revealed evenness (and denitrification) were significantly higher in *Juncus* than in *Spartina,* and highest in *Juncus*, 0-2 cm (Figure 3). Given that only evenness was significantly different when comparing the sediment samples, oscillations in evenness of particular community members likely resulted in the differences observed in the communities when ordinated (Figure 4). These finding leads to the question: what is the importance of microbial diversity, and in particular, evenness, in terms of nutrient removal via denitrification particularly as marsh ecosystems face future changing conditions? In this regard Wittebolle et al. (2009) used denitrifying microbial microcosms to show that initial microbial community evenness favors functional stability. In these experiments richness and evenness were manipulated in these denitrifying microcosms and then subjected to salinity stress. These experiments revealed that initial evenness resulted in stability of the net ecosystem denitrification when faced with stress. This suggests that biodiversity protects important ecosystem functions when faced with stress e.g. the “insurance hypothesis” (Yachi and Loreau, 1999). Importantly, Wittebolle et al. (2009) showed that evenness, not richness, provides functional stability, both when faced with stress, and under no-stress conditions. Thus our findings of higher denitrification in the most even microbial communities (*Juncus*) are consistent with Wittebolle et al. (2009) and suggested that the plant-microbe interactions that lead to more even communities may provide insurance regarding the marsh ecosystem service of nutrient filtering (denitrification) when faced with changing environmental conditions e.g. sea-level rise and increasing salinity.

Our findings that microbial evenness is higher in *Juncus* than in *Spartina* and the potential implications in terms of function (denitrification) and stability, led to further analyses of the core (Turnbaugh *et al.*, 2007; Hamady and Knight, 2009), or stable microbial community. The core microbiome is thought to be critical to the functioning of a community in a particular environment, and may thus be used as a community health diagnostic and provide predictive capabilities regarding community response to environmental change (Shade and Handelsman, 2012). In our samples 21 ASVs were defined as core (20 occurred in 95% and all occurred in 90% of the samples). Of those 20 core members, only a Planctomycetes, most similar to the cultured *Rubinisphaera brasiliensis* (Schlesner, 1989), which, as discussed above, is a known nitrate reducing microorganism (Schlesner, 1989; Scheuner *et al.*, 2014), was positively correlated with denitrification rates.

The role of Planctomycetes in the nitrogen cycle is well documented, with anaerobic ammonium-oxidizing (anammox) bacteria recognized early on as members of this clade (Strous *et al.*, 1999) that were shown to be both present and active in the marine environment (Thamdrup and Dalsgaard, 2002; Kuypers *et al.*, 2003). However, bin 134’s genome lacked a hydrazine oxidase enzyme, which acts on hydrazine converting it to nitrogen gas during the anammox reaction (Kartal *et al.*, 2011). Bin 134’s genome does encode a hydroxylamine reductase, an enzyme participating in anammox; however, bin 134’s enzyme was annotated as a hybrid-cluster protein (hcp) important in protecting cells from nitrosative stress (J. Wang *et al.*, 2016). The unique occurrence of hcp in the MAG compared to the other *Pirellulaceae* discussed previously does not appear to be an assembly artifact as the GC content of this gene was 61%, which was the GC content for the entire genome, for the encoded 462 long amino acid. More recently this enzyme’s role has been revisited and shown to act as a high affinity nitric oxide reductase that evolves nitrous oxide in *Escherichia coli* (J. Wang *et al.*, 2016). If this enzyme serves the same function in Planctomycetes bin 134 then a complete denitrification pathway is encoded in this microbe (Figure 6), which helps reconcile the unexpected positive correlation the core Planctomycetes ASV had with denitrification rates. Taken together, a lack of a hydrazine oxidase in bin 134’s genome along with the annotation of its hydroxylamine reductase as an hcp suggested that salt marsh Planctomycetes represented by this MAG may not be anammox bacteria; however, the genome was not complete so this cannot be definitively ruled out.

The contribution of Planctomycetes to the nitrogen cycle was expanded beyond anammox to include dissimilatory nitrate reduction to ammonium (DNRA) (Kartal *et al.,* 2007). Although bin 134’s genome codes for nitrite reduction this gene was annotated as belonging to the *nir* gene family, not as a periplasmic cytochrome *c* nitrite reductase (Nrf) that is a necessary enzyme in DNRA (Damashek *et al.,* 2018). Although the lack of *nrf* genes in bin 134 may be the result of an incomplete genome, it could also suggest that this Planctomycetes carries out complete denitrification, as discussed above, rather than DNRA.

In terms of denitrification and plant type, the higher rates of denitrification in *Juncus* compared to *Spartina* dominated sediments, despite similar porewater nitrate concentrations, that we report herein are consistent with Elsey-Quirk *et al.* (2011) who reported 46% higher N turnover in *Juncus* than in *Spartina,* highlighting the importance of plant species on belowground processes. These two plant types have several functional differences, with *Spartina* being more tolerant of flooding, anoxic soils, and high salinities (Mendelssohn and Morris, 2000), while *Juncus* is less stress tolerant (Eleuterius, 1976; Wiegart and Freeman, 1990). Further, hydrogen sulfide concentrations in the *Juncus* sediments were lower than that in the *Spartina* patches (Figure 1B, inset), a pattern consistent with the findings of Ledford et al. (in review) where they measured six-fold higher porewater concentrations in the *Spartina* plots compared to *Juncus* plots. The relationship between higher denitrification rates but lower hydrogen sulfide concentrations in *Juncus* compared to *Spartina* is in agreement with prior investigation on the inhibitory effects of hydrogen sulfide on nitrification (Joye and Hollibaugh, 1995) and denitrification (Sorensen *et al.,* 1980) and greater transport of oxygen to the subsurface by the more productive *Juncus* marshes (Koretsky *et al.,* 2008). Koretsky et al. (2008) also measured higher porewater phosphate concentrations in a *Juncus* compared to the *Spartina* marsh and suggested that this difference was because of the higher productivity and greater turnover of belowground material in the *Juncus* marsh. Consistent with Koretsky’s assertion, in a study from our marsh site, Starr et al. (2018) reported that *Juncus* had higher productivity than *Spartina.* While Wilson et al. (2015) measured elevated hydrogen sulfide concentrations in *Spartina* patches, Miley & Kiene (2004) working in a nearby *Juncus* marsh found that despite some of the highest rates of sulfate reduction measured, pore water hydrogen sulfide concentrations were extremely low and suggested that hydrogen sulfide either precipitated in the presence of the highly abundant oxidized forms of iron to produce pyrite or was being rapidly reoxidized to sulfate, an interpretation that is consistent with the findings of Dollhopf et al. (2005). Regardless of the mechanisms, it appears that in sediments dominated by different plants at the same elevation *Juncus* patches are associated with more even microbial communities and that within these communities only a Planctomycetes was positively correlated with denitrification rates. Further, we also showed that an uncultured Planctomycetes with a genome assembled from metagenome sequence data may encode a complete denitrification pathway resolving why a Planctomycetes was the only microbe positively correlated with denitrification rates. Thus, our results highlight how plant-microbe interactions influence marsh ecosystems function (i.e. denitrification).

## Conclusion

This study provided new insights into how plant-microbe interactions influence microbial community structure, particularly evenness, with more even communities leading to enhanced denitrification. Further, the analysis of a genome from an uncultured Planctomycetes provided new resolution on the role of this clade in nitrogen cycling in a salt marsh. This new understanding is important given that worldwide marshes are disappearing at an alarming rate (Zedler and Kercher, 2005), thereby reducing the critical ecosystem services they provide such as nutrient retention and removal (Costanza *et al.,* 1997). Wetlands remove 1% of the global fixed nitrogen (Seitzinger *et al.,* 2006) with salt marshes removing, on average, a remarkable 33% of the nitrogen input they receive through denitrification (Jordan *et al.,* 2011). Factors such as rising seas, lack of sediment input, eutrophication, urbanization, and diking and channeling (Zedler and Kercher, 2005) are affecting marshes and by changing the composition of the vegetation and their associated microbial community will alter the ecosystem functions they provide.

## Acknowledgments

We would like to thank A. Kleinhuizen and L. Linn with assistance in the laboratory. We would also like to thank Philip Hugenholtz and Gene Tyson for the opportunity to analyze metagenomic data at the Australian Centre for Ecogenomics. This project was funded by the National Science Foundation’s Division of Chemical, Bioengineering, Environmental and Transport Systems grants 1438092 and 1643486. Metagenomic sequencing was provided by the Joint Genome Institute through a small-scale community sequencing project grant 503678.

## Author Contributions Statement

BM and PC collected samples and determined denitrification rates. LK carried out DNA extractions, library preparation and 16S rRNA gene sequencing. OUM carried out bioinformatics and statistical analyses of the 16S rRNA gene sequence data. JZ and OUM carried metagenome assembly and analyzed MAGs. BM and OUM wrote the manuscript.

## References cited

Adams, J.M., Faure, H., Faure-Denard, L., McGlade, J.M., and Woodward, F.I. (1990) Increases in terrestrial carbon storage from the Last Glacial Maximum to the present. Nature 348: 711–714.

Anderson, I., Tobias, C., Neikirk, B., and Wetzel, R.L. (1997) Development of a process-based nitrogen mass balance model for a Virginia (USA) Spartina alterniflora salt marsh: implications for net DIN flux. Mar. Ecol. Prog. Ser. 159: 13–27.

Angell, J.H., Peng, X., Ji, Q., Craick, I., Jayakumar, A., Kearns, P., Ward, B.B., and Bowen, J.L. (2018) Community Composition of Nitrous Oxide-Related Genes in Salt Marsh Sediments Exposed to Nitrogen Enrichment. Front. Microbiol. 9: 1–13.

Angermeyer, A., Crosby, S.C., and Huber, J.A. (2018) Salt marsh sediment bacterial communities maintain original population structure after transplantation across a latitudinal gradient. PeerJ 1–21.

Apprill, A., McNally, S., Parsons, R., and Weber, L. (2015) Minor revision to V4 region SSU rRNA 806R gene primer greatly increases detection of SAR11 bacterioplankton. Aquat. Microb. Ecol. 75: 129–137.

Battaglia, L.L., Woodrey, M.S., Peterson, M.S., Dillon, K.S., and Visser, J.M. (2012) Wetland Ecosystems of the Northern Gulf Coast in Wetland Habitats of North America: Ecology and Conservation Concerns. In, Batzer,D.P. and Baldwin,A.H. (eds), Wetland Habitats of North America: Ecology and Conservation Concerns.,pp. 75–88.

Bowen, J.L., Kearns, P.J., Byrnes, J.E.K., Wigginton, S., Allen, W.J., Greenwood, M., Tran, K., Yu, J., Cronin, J.T., and Meyerson, L.A. (2017) Lineage overwhelms environmental conditions in determining rhizosphere bacterial community structure in a cosmopolitan invasive plant. Nat. Commun. 8: 433.

Bowen, J.L., Morrison, H.G., Hobbie, J.E., and Sogin, M.L. (2012) Salt marsh sediment diversity: a test of the variability of the rare biosphere among environmental replicates. ISME J 6: 2014–2023.

Bridgham, S.D., Megonigal, J.P., Keller, J.K., Bliss, N.B., and Trettin, C. (2006) The carbon balance of North American wetlands. Wetlands 26: 889–916.

Burke, D.J., Hamerlynck, E.P., and Hahn, D. (2002) Interactions among Plant Species and Microorganisms in Salt Marsh Sediments. Appl. Environ. Microbiol. 68: 1157–1164.

Callahan, B.J., Mcmurdie, P.J., and Holmes, S.P. (2017) Exact sequence variants should replace operational taxonomic units in marker-gene data analysis. ISME J. 11: 2639–2643.

Callahan, B.J., Mcmurdie, P.J., Rosen, M.J., Han, A.W., Johnson, A.J.A., and Holmes, S.P. (2016) DADA2: High-resolution sample inference from Illumina amplicon data. Nat. Methods 13: 581–583.

Cao, Y.P., Green, P.G., and Holden, P.A. (2008) Microbial community composition and denitrifying enzyme activities in salt marsh sediments. Appl Env. Microbiol 74: 7585–7595.

Caporaso, J.G., Kuczynski, J., Stombaugh, J., Bittinger, K., Bushman, F., Costello, E.K., Fierer, N., Gonzalez, A., Goodrich, J.K., Gordon, J.I., Huttley, G.A., Kelley, S.T., Knights, D., Koenig, J.E., Ley, R.E., Lozupone, C.A., McDonald, D., Muegge, B.D., Pirrung, M., et al. (2010) QIIME allows analysis of high-throughput community sequencing data. Nat. Methods 7: 335–336.

Caporaso, J.G., Lauber, C.L., Walters, W. a, Berg-Lyons, D., Lozupone, C. a, Turnbaugh, P.J., Fierer, N., and Knight, R. (2011) Global patterns of 16S rRNA diversity at a depth of millions of sequences per sample. Proc. Natl. Acad. Sci. U. S. A. 108 Suppl: 4516–22.

Caporaso, J.G., Lauber, C.L., Walters, W.A., Berg-Lyons, D., Huntley, J., Fierer, N., Owens, S.M., Betley, J., Fraser, L., Bauer, M., Gormley, N., Gilbert, J.A., Smith, G., and Knight, R. (2012) Ultra-high-throughput microbial community analysis on the Illumina HiSeq and MiSeq platforms. ISME J. 6: 1621–4.

Chao, A. (1984) Nonparametric Estimation of the Number of Classes in a Population. Scand. J. Stat. 11: 265–270.

Chaumeil, P.-A., Mussig, A.J., Hugenholtz, P., and Parks, D.H. (2019) GTDB-Tk: a toolkit to classify genomes with the Genome Taxonomy Database. Bioinformatics.

Chen, I.-M.A., Chu, K., Palaniappan, K., Pillay, M., Ratner, A., Huang, J., Huntemann, M., Varghese, N., White, J.R., Seshadri, R., Smirnova, T., Kirton, E., Jungbluth, S.P., Woyke, T., Eloe-Fadrosh, E.A., Ivanova, N.N., and Kyrpides, N.C. (2019) IMG/M v.5.0: an integrated data management and comparative analysis system for microbial genomes and microbiomes. Nucleic Acids Res. 47: D666–D677.

Chmura, G.L., Anisfeld, S.C., Cahoon, D.R., and Lynch, J.C. (2003) Global carbon sequestration in tidal, saline wetland soils. Global Biogeochem. Cycles 17: 22.

Cleary, D.F.R., Coelho, F.J.R.C., Oliveira, V., Gomes, N.C.M., and Polònia, A.R.M. (2017) Sediment depth and habitat as predictors of the diversity and composition of sediment bacterial communities in an inter-tidal estuarine environment. Mar. Ecol. 38: 1–15.

Cleary, D.F.R., Polónia, A.R.M., Sousa, A.I., LillebØ, A.I., Queiroga, H., and Gomes, N.C.M. (2016) Temporal dynamics of sediment bacterial communities in monospecific stands of Juncus maritimus and Spartina maritima. Plant Biol. 18: 824–834.

Costanza, R., d’Arge, R., de Groot, R., Farber, S., Grasso, M., Hannon, B., Limburg, K., Naeem, S., O’Neill, R. V, Paruelo, J., Raskin, R.G., Sutton, P., and van den Belt, M. (1997) The value of the world’s ecosystem services and natural capital. Nature 387: 253–260.

Costanza, R., Groot, R. De, Sutton, P., Ploeg, S. Van Der, Anderson, S.J., Kubiszewski, I., Farber, S., and Turner, R.K. (2014) Changes in the global value of ecosystem services. Glob. Environ. Chang. 26: 152–158.

Damashek, J., Francis, C.A., and Francis, C.A. (2018) Microbial Nitrogen Cycling in Estuaries: From Genes to Ecosystem Processes. Estuaries and Coasts 41: 626–660.

Davis, D.A., Gamble, M.D., Bagwell, C.E., Bergholz, P.W., and Lovell, C.R. (2011) Responses of salt marsh plant rhizosphere diazotroph assemblages to changes in marsh elevation, edaphic conditions and plant host species. Microb. Ecol. 61: 386–398.

DeLaune, R.D., Baumann, R.H., and Gosselink, J.G. (1983) Relationships among vertical accretion, coastal submergence, and erosion in a Louisiana Gulf Coast marsh. J. Sediment. Res. 53: 147–157.

Dixon, R.K. and Krankina, O. (1995) Can the terrestrial biosphere be managed to conserve and sequester carbon? In, Beran,M.A. (ed), Carbon Sequestration in the Biosphere. Springer-Verlag, Netherlands, pp. 153 – 179.

Dollhopf, S.L., Hyun, J.-H., Smith, A.C., Adams, H.J., O’Brien, S., and Kostka, J.E. (2005) Quantification of ammonia-oxidizing bacteria and factors controlling nitrification in salt marsh sediments. Appl Env. Microbiol 71: 240–246.

Duarte, C., Losada, I., Hendriks, I., Mazarrasa, I., and Marba, N. (2013) The role of coastal plant communities for climate change mitigation and adaptation. Nat. Clim. Chang. 3: 961–968.

Eleuterius, L.N. (1976) The Distribution of Juncus roemerianus in the Salt Marshes of North America. Chesap. Sci. 17: 289–292.

Elsey-Quirk, T., Seliskar, D.M., and Gallagher, J.L. (2011) Nitrogen Pools of Macrophyte Species in a Coastal Lagoon Salt Marsh: Implications for Seasonal Storage and Dispersal. Estuaries and Coasts 34: 470–482.

Elsey-Quirk, T., Seliskar, D.M., Sommerfield, C.K., and Gallagher, J.L. (2011) Salt marsh carbon pool distribution in a mid-Atlantic lagoon, USA: sea level rise implications. Wetlands 31: 87–99.

Elsey-Quirk, T. and Unger, V. (2018) Geomorphic influences on the contribution of vegetation to soil C accumulation and accretion in Spartina alterniflora marshes. Biogeosciences 15: 379–397.

Fourqurean, J.W., Duarte, C.M., Kennedy, H., Marbà, N., Holmer, M., Mateo, M.A., Apostolaki, E.T., Kendrick, G.A., and Krause-jensen, D. (2012) Seagrass ecosystems as a globally significant carbon stock. Nat. Geosci. 5: 505–509.

Gallagher, J.L. and Plumley, F.G. (1979) Underground Biomass Profiles and Productivity in Atlantic Coastal Marshes. Am. J. Bot. 66: 156–161.

Gillies, L.E., Thrash, J.C., de Rada, S., Rabalais, N.N., and Mason, O.U. (2015) Archaeal enrichment in the hypoxic zone in the northern Gulf of Mexico. Environ. Microbiol. 17: 3847–3856.

Glöckner, F.O., Kube, M., Bauer, M., Teeling, H., Lombardot, T., Ludwig, W., Gade, D., Beck, A., Borzym, K., Heitmann, K., Rabus, R., Schlesner, H., Amann, R., and Reinhardt, R. (2003) Complete genome sequence of the marine planctomycete Pirellula sp. strain 1. Proc. Natl. Acad. Sci. 100: 8298 LP–8303.

Gribsholt, B., Kostka, J.E., and Kristensen, E. (2003) Impact of fiddler crabs and plant roots on sediment biogeochemistry in a Georgia saltmarsh. Mar. Ecol. Prog. Ser. 259: 237–251.

Hamady, M. and Knight, R. (2009) Microbial community profiling for human microbiome projects: tools, techniques, and challenges. Genome Res. 1141–1152.

Hamersley, M.R. and Howes, B.L. (2005) Coupled nitrification-denitrification measured in situ in a Spartina alterniflora marsh with a 15NH4+ tracer. Mar. Ecol. Prog. Ser. 299: 123–135.

Hopkinson, C.S. and Giblin, A.E. (2008) Nitrogen Dynamics of Coastal Salt Marshes. In, Capone,D.G., Bronk,D.A., Mulholland,M.R., and Carpenter,E.J. (eds), Nitrogen in the marine environment. Academic Press, San Diego, pp. 991–1036.

Howard, J., Sutton-Grier, A., Herr, D., Kleypas, J., Landis, E., Mcleod, E., Pidgeon, E., and Simpson, S. (2017) Clarifying the role of coastal and marine systems in climate mitigation. Front. Ecol. Environ. 15: 42–50.

Jordan, S.J., Stoffer, J., and Nestlerode, J.A. (2011) Wetlands as Sinks for Reactive Nitrogen at Continental and Global Scales: A Meta-Analysis. Ecosystems 14: 144–155.

Joye, S.B. and Hollibaugh, T. (1995) Influence of sulfide inhibition of nitrification on nitrogen regeneration in sediments. 270: 623–625.

Kang, D.D., Froula, J., Egan, R., and Wang, Z. (2015) MetaBAT, an efficient tool for accurately reconstructing single genomes from complex microbial communities. PeerJ 3: e1165.

Kang, D.D., Li, F., Kirton, E., Thomas, A., Egan, R., An, H., and Wang, Z. (2019) MetaBAT 2: an adaptive binning algorithm for robust and efficient genome reconstruction from metagenome assemblies. PeerJ 7: e7359.

Kartal, B., Keltjens, J., and Jetten, M. (2011) Metabolism and Genomics of Anammox Bacteria. In, Ward,B., Arp,D., and Klotz,M. (eds), Nitrification. ASM Press, Washington DC, pp. 181–200.

Kartal, B., Kuypers, M.M.M., Lavik, G., Schalk, J., op den Camp, H.J.M., Jetten, M.S.M., and Strous, M. (2007) Anammox bacteria disguised as denitrifiers: nitrate reduction to dinitrogen gas via nitrite and ammonium. Environ. Microbiol. 9: 635–642.

King, G.M., Klug, M.J., Wiegert, R.G., and Chalmers, A.G. (1982) Relation of Soil Water Movement and Sulfide Concentration to Spartina alterniflora Production in a Georgia Salt Marsh. Science 218: 61–63.

Koop-Jakobsen, K. and Giblin, A.E. (2010) The effect of increased nitrate loading on nitrate reduction via denitrification and DNRA in salt marsh sediments. Limnol. Oceanogr. 55: 789–802.

Koop-Jakobsen, K. and Wenzhöfer, F. (2015) The Dynamics of Plant-Mediated Sediment Oxygenation in Spartina anglica Rhizospheres—a Planar Optode Study. Estuaries and Coasts 38: 1–13.

Koretsky, C.M., Haveman, M., Cuellar, A., Beuving, L., Shattuck, T., and Wagner, M. (2008) Influence of Spartina and Juncus on Saltmarsh Sediments. I. Pore Water Geochemistry. Chem. Geol. 255: 87–99.

Kostka, J.E., Gribsholt, B., Petrie, E., Dalton, D., Skelton, H., and Kristensen, E. (2002) The rates and pathways of carbon oxidation in bioturbated saltmarsh sediments. Limnol. Oceanogr. 47: 230–240.

Kuypers, M.M.M., Sliekers, A.O., Lavik, G., Schmid, M., Jørgensen, B.B., Kuenen, J.G., Sinninghe Damsté, J.S., Strous, M., and Jetten, M.S.M. (2003) Anaerobic ammonium oxidation by anammox bacteria in the Black Sea. Nature 422: 608–611.

Langley, J.A. and Megonigal, J.P. (2010) Ecosystem response to elevated CO2 levels limited by nitrogen-induced plant species shift. Nature 466: 96–99.

Linthurst, R.A. and Seneca, E.D. (1980) The effects of standing water and drainage potential on the Spartina Alterniflora-substrate complex in a North Carolina salt marsh. Estuar. Coast. Mar. Sci. 11: 41–52.

Mcleod, E., Chmura, G.L., Bouillon, S., Salm, R., Björk, M., Duarte, C.M., Lovelock, C.E., Schlesinger, W.H., and Silliman, B.R. (2011) A blueprint for blue carbon: toward an improved understanding of the role of vegetated coastal habitats in sequestering CO2. Front. Ecol. Environ. 9: 552–560.

Mendelssohn, I.A. and Morris, J.T. (2000) Eco-physiological controls on the productivity of Spartina Alterniflora Loisel. In, D.A. Kreeger (ed), Concepts and Controversies in Tidal Marsh Ecology. Kluwer Academic, Dordrecht.

Meng, J., Xu, J., Qin, D., He, Y., Xiao, X., and Wang, F. (2014) Genetic and functional properties of uncultivated MCG archaea assessed by metagenome and gene expression analyses. ISME J. 8: 650–659.

Miley, G.A. and Kiene, R.P. (2004) Sulfate reduction and porewater chemistry in a gulf coastJuncus roemerianus(Needlerush) marsh. Estuaries and Coasts 27: 472–481.

Mitsch, W.J. and Gosselink, J.G. (2015) Wetlands 5th ed. John Wiley & Sons, Inc., Hoboken, NJ, NJ.

Moffett, K.B. and Gorelick, S.M. (2016) Relating salt marsh pore water geochemistry patterns to vegetation zones and hydrologic influences. WATER Resour. Res. 52: 1729–1745.

Neubauer, S.C. (2013) Ecosystem Responses of a Tidal Freshwater Marsh Experiencing Saltwater Intrusion and Altered Hydrology. Estuaries and Coasts 36: 491–507.

Newell, S.Y. (2001) Multiyear patterns of fungal biomass dynamics and productivity within naturally decaying smooth cordgrass shoots. Limnol. Oceanogr. 46: 573–583.

Nyman, J.A., Walters, R.J., Delaune, R.D., and Patrick, W.H. (2006) Marsh vertical accretion via vegetative growth. Estuar. Coast. Shelf Sci. 69: 370–380.

Odum, E. (1959) Fundamentals of Ecology WB Saunders Comp., Philadelphia.

Oliveira, V., Santos, A.L., Aguiar, C., Santos, L., Salvador, Â.C., Gomes, N.C.M., Silva, H., Rocha, S.M., Almeida, A., and Cunha, Â. (2012) Prokaryotes in salt marsh sediments of Ria de Aveiro: Effects of halophyte vegetation on abundance and diversity. Estuar. Coast. Shelf Sci. 110: 61–68.

Oliveira, V., Santos, A.L., Coelho, F., Gomes, N.C.M., Silva, H., Almeida, A., and Cunha, Â. (2010) Effects of Monospecific Banks of Salt Marsh Vegetation on Sediment Bacterial Communities. Microb. Ecol. 60: 167–179.

Parada, A.E., Needham, D.M., and Fuhrman, J.A. (2015) Every base matters: Assessing small subunit rRNA primers for marine microbiomes with mock communities, time series and global field samples. Environ. Microbiol. 18: 1403–1414.

Parks, D.H., Chuvochina, M., Waite, D.W., Rinke, C., Skarshewski, A., Chaumeil, P.-A., and Hugenholtz, P. (2018) A standardized bacterial taxonomy based on genome phylogeny substantially revises the tree of life. Nat. Biotechnol. 36: 996–1004.

Parks, D.H., Imelfort, M., Skennerton, C.T., Hugenholtz, P., and Tyson, G.W. (2015) CheckM: assessing the quality of microbial genomes recovered from isolates, single cells, and metagenomes. Genome Res. 25: 1043–1055.

Paulson, J.N., Stine, O.C., Bravo, H.C., and Pop, M. (2013) Differential abundance analysis for microbial marker-gene surveys. Nat. Methods 10: 1200–2.

Pielou, E.C. (1966) The measurement of diversity in different types of biological collections. J. Theor. Biol. 13: 131–144.

Pomeroy, L.R. and Wiegert, R.G. (1981) Ecology of a salt marsh Springer-Verlag, New York.

Revelle, W. (2018) psych: Procedures for Personality and Psychological Research. https://CRAN.R-project.org/package=psych.

Rietl, A.J., Overlander, M.E., Nyman, A.J., and Jackson, C.R. (2016) Microbial Community Composition and Extracellular Enzyme Activities Associated with Juncus roemerianus and Spartina alterniflora Vegetated Sediments in Louisiana Saltmarshes. Microb. Ecol. 71: 290–303.

Sahagian, D. and Melack, J. (1998) Global wetland distribution and functional characterization: Trace gases and the hydrologic cycle Intl. Geosphere Biosphere Programme Secretariat.

Scheuner, C., Tindall, B.J., Lu, M., Nolan, M., Lapidus, A., Cheng, J., Goodwin, L., Pitluck, S., Huntemann, M., Liolios, K., Pagani, I., Mavromatis, K., Ivanova, N., Pati, A., Chen, A., Palaniappan, K., Jeffries, C.D., Hauser, L., Land, M., et al. (2014) Complete genome sequence of Planctomyces brasiliensis type strain (DSM 5305(T)), phylogenomic analysis and reclassification of Planctomycetes including the descriptions of Gimesia gen. nov., Planctopirus gen. nov. and Rubinisphaera gen. nov. and emended d. Stand Genomic Sci 8: 9–10.

Schlesner, H. (1989) Planctomyces brasiliensis sp. nov., a Halotolerant Bacterium from a Salt Pit. Syst. Appl. Microbiol. 12: 159–161.

Schubauer, J.P. and Hopkinson, C.S. (1984) Above- and belowground emergent macrophyte production and turnover in a coastal marsh ecosystem, Georgia. Limnol. Oceanogr. 29: 1052–1065.

Seitzinger, S., Harrison, J.A., Böhlke, J.K., Bouwman, A.F., Lowrance, R., Peterson, B., Tobias, C., and Drecht, G. Van (2006) Denitrification across landscapes and waterscapes: a synthesis. Biogeosciences Discuss 16: 2064–2090.

Shade, A. and Handelsman, J. (2012) Beyond the Venn diagram: The hunt for a core microbiome. Environ. Microbiol. 14: 4–12.

Shannon, C.E. and Weaver, W. (1949) The mathematical theory of communication The University of Illinois Press, Urbana.

Solomon, S., Qin, D., Manning, M., Chen, Z., Marquis, M., Averyt, K.B., Tignor, M., and Miller, H.L. (2007) Contribution of Working Group I to the Fourth Assessment Report of the Intergovernmental Panel on Climate Change Cambridge University Press, Cambridge, United Kingdom and New York, NY, USA.

Sorensen, J. (1978) Denitrification rates measured in a marine sediment as measured by acetlyene inhibition technique. Appl. Environ. Microbiol. 36: 139–143.

Sorensen, J., Tiedje, J.M., and Firestone, R.B. (1980) Inhibition by Sulfide of Nitric and Nitrous Oxide reduction by denitrifying Pseudomonas fluorescens. Appl. Environ. Microbiol. 39: 105–108.

Starr, G., Staudhammer, C.L., Starr, G., Jarnigan, J.R., Staudhammer, C.L., and Cherry, J.A. (2018) Variation in ecosystem carbon dynamics of saltwater marshes in the northern Gulf of. Wetl. Ecol. Manag. 26: 581–596.

Stout, J. (1984) The Ecology of Irregularly Flooded Salt Marshes of the Northeastern Gulf of Mexico: a Community Profile U.S. Department of the Interior, Fish and Wildlife Service, Washington DC.

Strous, M., Fuerst, J.A., Kramer, E.H.M., Logemann, S., Muyzer, G., van de Pas-Schoonen, K.T., Webb, R., Kuenen, J.G., and Jetten, M.S.M. (1999) Missing lithotroph identified as new planctomycete. Nature 400: 446–449.

Sunagawa, S., Coelho, L.P., Chaffron, S., Kultima, J.R., Labadie, K., Salazar, G., Djahanschiri, B., Zeller, G., Mende, D.R., Alberti, A., Cornejo-castillo, F.M., Costea, P.I., Cruaud, C., Ovidio, F., Engelen, S., Ferrera, I., Gasol, J.M., Guidi, L., Hildebrand, F., et al. (2015) Structure and function of the global ocean microbiome. 348: 1–10.

Thamdrup, B. and Dalsgaard, T. (2002) Production of N2 through anaerobic ammonium oxidation coupled to nitrate reduction in marine sediments. Appl Env. Microbiol 68: 1312–1318.

Tiner, R.W. (1984) Wetlands of the United States--Current status and recent trends. U.S. Fish and Wildlife Service Report, Washington, D.C., USA.

Tobias, C.R., Macko, S.A., Anderson, I.C., Canuel, E.A., and Harvey, J.W. (2001) Tracking the fate of a high concentration groundwater nitrate plume through a fringing marsh: A combined groundwater tracer and in situ isotope enrichment study. Limnol. Oceanogr. 46: 1977–1989.

Turnbaugh, P.J., Ley, R.E., Hamady, M., Fraser-Liggett, C.M., Knight, R., and Gordon, J.I. (2007) The Human Microbiome Project. Nature 449: 804–810.

Valiela, I. and Cole, M.L. (2002) Comparative evidence that salt marshes and mangroves may protect seagrass meadows from land-derived nitrogen loads. Ecosystems 5: 92–102.

Valiela, I., Geist, M., McClelland, J., and Tomasky, G. (2000) Nitrogen loading from watersheds to estuaries: Verification of the Waquoit Bay Nitrogen Loading Model. Biogeochemistry 49: 277–293.

Valiela, I. and Teal, J.M. (1979) The nitrogen budget of a salt marsh ecosystem. Nature 280: 652–656.

Valiela, I., Teal, J.M., and Persson, N.Y. (1976) Production and dynamics of experimentally enriched salt marsh vegetation: Belowground biomass1. Limnol. Oceanogr. 21: 245–252.

Velinsky, D.J., Paudel, B., Quirk, T., Piehler, M., and Smyth, A. (2017) Salt Marsh Denitrification Provides a Significant Nitrogen Sink in Barnegat Bay, New Jersey. J. Coast. Res. 78: 70–78.

Wang, J., Vine, C.E., Balasiny, B.K., Rizk, J., Bradley, C.L., Tinajero-Trejo, M., Poole, R.K., Bergaust, L.L., Bakken, L.R., and Cole, J.A. (2016) The roles of the hybrid cluster protein, Hcp and its reductase, Hcr, in high affinity nitric oxide reduction that protects anaerobic cultures of Escherichia coli against nitrosative stress. Mol. Microbiol. 100: 877–892.

Wang, M., Yang, P., Salles, J.F., and Saleem, M. (2016) Distribution of Root-Associated Bacterial Communities Along a Salt-Marsh Primary Succession. Front Plant Sci 6: 1188.

Ward, L.G., Zaprowski, B.J., Trainer, K.D., and Davis, P.T. (2008) Stratigraphy, pollen history and geochronology of tidal marshes in a Gulf of Maine estuarine system: Climatic and relative sea level impacts. Mar. Geol. 256: 1–17.

Watson, R.T., Noble, I.R., Bolin, B., Ravindranath, N.H., Verardo, D.J., and Dokken, D.J. (2000) Land use, land-use change, and forestry. In, Intergovernmental Panel on Climate Change Special Report. Cambridge University Press, New York.

Wiegart, R.G. and Freeman, B.J. (1990) Tidal marshes of the southeastern Atlantic coast: a community profile. U.S. Fish Wildl. Serv. 85: 1–80.

Wiegert, R.G., Chalmers, A.G., and Randerson, P.F. (1983) Productivity Gradients in Salt Marshes: The Response of Spartina Alterniflora to Experimentally Manipulated Soil Water Movement. Oikos 41: 1–6.

Wilson, B.J., Mortazavi, B., and Kiene, R.P. (2015) Spatial and temporal variability in carbon dioxide and methane exchange at three coastal marshes along a salinity gradient in a northern Gulf of Mexico estuary. Biogeochemistry 123: 329–347.

Wittebolle, L., Marzorati, M., Clement, L., Balloi, A., Daffonchio, D., Heylen, K., De Vos, P., Verstraete, W., and Boon, N. (2009) Initial community evenness favours functionality under selective stress. Nature 458: 623–626.

Wu, Y.-. W., Tang, Y.-. H., Tringe, S.G., Simmons, B.A., and Singer, S.W. (2014) MaxBin: an automated binning method to recover individual genomes from metagenomes using an expectation-maximization algorithm. Microbiome 2: 26.

Yachi, S. and Loreau, M. (1999) Biodiversity and ecosystem productivity in a fluctuating environment: The insurance hypothesis. Proc. Natl. Acad. Sci. 96: 1463–1468.

Yilmaz, P., Parfrey, L.W., Yarza, P., Gerken, J., Pruesse, E., Quast, C., Schweer, T., Peplies, J., Ludwig, W., and Glockner, F.O. (2014) The SILVA and “All-species Living Tree Project (LTP)” taxonomic frameworks. Nucleic Acids Res. 42: D643–D648.

Zedler, J.B. and Kercher, S. (2005) WETLAND RESOURCES: Status, Trends, Ecosystem Services, and Restorability. Annu. Rev. Environ. Resour. 30: 39–74.

